# Dual Hypocretin Receptor Antagonism Reduces Oxycodone Seeking During Abstinence

**DOI:** 10.1101/2025.04.05.647321

**Authors:** Kyle R. Samson, Alison R. Bashford, Rodrigo A. España

**Author notes:** **Corresponding Author** Rodrigo A. España, Ph.D., Department of Neurobiology and Anatomy, Drexel University College of Medicine, 2900 W Queen Ln, Philadelphia, PA 19129.

## Abstract

A major barrier in the treatment of opioid use disorder is persistent drug craving during abstinence. While opioid-based medications have been used to treat opioid use disorder for decades, there is an urgent need for novel, non-opioid-based pharmacotherapies. The hypocretin/orexin (hypocretin) system is a promising target for treating opioid use disorder due to its influence on motivation for drugs of abuse through actions on dopamine transmission. We recently showed that intermittent access (IntA) to oxycodone promoted sustained oxycodone seeking and alterations in dopamine transmission during abstinence. In the current studies, we investigated to what extent suvorexant, an FDA-approved dual hypocretin receptor antagonist, reduces oxycodone seeking and restores dopamine function during abstinence. Results indicated that IntA to oxycodone produced sustained cue-induced oxycodone seeking after a 14-day abstinence period, which was associated with reduced dopamine uptake in the nucleus accumbens core as we have previously shown. Treatment with suvorexant 24 h prior to a cue-induced seeking test significantly reduced oxycodone seeking and normalized aberrant dopamine uptake. These findings suggest that targeting hypocretin receptors may be a promising strategy for reducing opioid craving and associated neuroadaptations, thus lowering the risk of relapse.

## Introduction

Individuals with opioid use disorder experience intense and persistent drug craving during periods of abstinence, which promotes relapse and significantly contributes to opioid overdoses (Vafaie and Kober, 2022). Our previous report in rats demonstrated that self-administration of oxycodone on an intermittent access (IntA) schedule promoted sustained oxycodone seeking following abstinence, along with reduced dopamine uptake rates in the nucleus accumbens core (Samson et al., 2022). This finding is consistent with studies in which humans and non-human primates show reduced dopamine transporter (DAT) availability in the striatum during opioid abstinence (Xiao et al., 2006; Yeh et al., 2012; Liu et al., 2013; Yuan et al., 2017). Together, these observations suggest that persistent opioid craving during abstinence is linked to reduced dopamine uptake rates, highlighting the potential of normalizing DAT function as a strategy to prevent relapse.

Approved pharmacotherapies for opioid use disorder are opioid-based, carry a significant risk of misuse [7], and have not effectively curbed the opioid crisis, which emphasizes the urgent need for novel, non-opioid-based treatments to reduce craving and relapse. The National Institute on Drug Abuse has named the hypocretin system as a top priority for developing treatments for opioid use disorder (Rasmussen et al., 2019), due to extensive evidence that hypocretin antagonists decrease behavioral and dopamine responses to opioids (Simmons et al., 2017; Brodnik et al., 2020). For example, antagonism of hypocretin receptor 1 (HcrtR1) reduced opioid self-administration (Smith and Aston-Jones, 2012; Matzeu and Martin-Fardon, 2020), motivation for opioids (Smith and Aston-Jones, 2012; Porter-Stransky et al., 2017; Fragale et al., 2019; Mohammadkhani et al., 2020; Fragale et al., 2021), and cue-induced reinstatement of opioid seeking (Smith and Aston-Jones, 2012; Matzeu and Martin-Fardon, 2020; Fragale et al., 2021). Fewer studies have explored the involvement of hypocretin receptor 2 (HcrtR2) in opioid-associated behavior with one study showing that HcrtR2 antagonism reduced heroin self-administration (Schmeichel et al., 2015), but others indicating that HcrtR2 antagonism had no effect on oxycodone self-administration or cued reinstatement of oxycodone seeking (Matzeu and Martin-Fardon, 2020).

A few recent studies have explored the effects of dual hypocretin receptor antagonism on opioid-seeking behavior. For instance, the FDA-approved dual hypocretin receptor antagonist, suvorexant reduced oxycodone self-administration and conditioned reinstatement of oxycodone seeking (Illenberger et al., 2023) and reduced motivation for fentanyl (O’Connor et al., 2020). Similarly, daily treatment with DORA-12 during oxycodone abstinence reduced conditioned reinstatement of oxycodone seeking (Illenberger et al., 2023).

Despite these observations, the extent to which hypocretin receptors influence opioid seeking and dopamine transmission during abstinence remains unclear. To address this gap, we investigated to what degree suvorexant reduces opioid seeking and normalizes dopamine transmission following abstinence from oxycodone. Rats were trained to self-administer oxycodone using an IntA schedule of reinforcement, followed by a 14-day abstinence period. To assess the effects of dual hypocretin receptor antagonism on oxycodone seeking, suvorexant or vehicle was administered during abstinence, one day before a cue-induced seeking test on abstinence day (AD)14. One day following the AD14 seeking test, we performed fast-scan cyclic voltammetry (FSCV) in the NAc core to assess changes in dopamine transmission. Additionally, to assess whether suvorexant modulated dopamine responses to oxycodone in the NAc core, we applied oxycodone to brain slices and measured evoked dopamine release (peak height) and dopamine uptake rate.

## Materials and Methods

### Animals

Adult female (210-260g) and male (325-430g) Long Evans rats (Envigo, Frederick, MD, USA) were maintained on a 12-hour reverse light/dark cycle (1500 lights on; 0300 lights off) with *ad libitum* access to food and water. After arrival, rats were given at least 7 days to acclimate to the animal facility prior to surgery. All protocols and animal procedures were conducted in accordance with National Institutes of Health Guide for the Care and Use of Laboratory Animals under supervision of the Institutional Animal Care and Use Committee at Drexel University College of Medicine.

### Drugs

Oxycodone hydrochloride was purchased from Spectrum Pharmacy Products (Spectrum Laboratory Products, Inc.). For self-administration experiments, oxycodone was dissolved in 0.9% physiological saline. For FSCV experiments, oxycodone was dissolved in artificial cerebrospinal fluid (aCSF). Suvorexant was provided by Dr. Yanan Zhang (Research Triangle Institute, NC, USA). Suvorexant (30 mg/kg) was dissolved in dimethyl sulfoxide (DMSO; 100%) at a fixed volume (100 nl) (Simmons et al., 2017; O’Connor et al., 2020).

### Intravenous catheter surgery

Rats were anesthetized using 2.5% isoflurane and implanted with a silastic catheter (ID, 0.012 in OD, 0.025 in. Access Technologies, Skokie, IL) into the right jugular vein for intravenous delivery of oxycodone. The catheter was connected to a cannula which exited through the skin on the dorsal surface in the region of the scapulae. Ketoprofen (Patterson Veterinary, Devens, MA; 5mg/kg s.c. of 5 mg/ml) and Enrofloxacin (Norbrook, Northern Ireland; 5 mg/kg s.c. of 5 mg/ml) were provided at the time of surgery and a second dose was given 12 h later. In addition, combined antibiotic and analgesic powder (Neo-Predef, Kalamazoo, MI) was applied around the chest and back incisions. Rats were subsequently singly housed and recovered for 5 days prior to self-administration. Intravenous catheters were manually flushed with Gentamicin (5 mg/kg i.v. of 5 mg/ml) in heparinized saline every day during recovery.

### Self-Administration

All measures of self-administration were recorded using custom created Ghost Software (Bernosky-Smith et al., 2016). Each rat underwent one oxycodone self-administration session per day from 10:00-16:00. Rats were first trained to self-administer oxycodone for 6 h under a fixed ratio 1 schedule whereby a single active lever press initiated an intravenous injection of oxycodone (0.1 mg/kg, infused over 2.5 s - 4.5 s) with a 20 s cue light presentation above the active lever and a 20 s timeout during which the levers retracted. This dose was chosen based on previous oxycodone self-administration studies (Thompson et al., 2000; Blackwood et al., 2019b; Blackwood et al., 2019a; Samson et al., 2022). Acquisition of the behavior occurred when a rat obtained ≥ 20 infusions and inactive lever presses were less than half the number of active lever presses in two consecutive sessions.

### Intermittent Access

Following acquisition, rats were switched to an IntA schedule of reinforcement (Zimmer et al., 2012), which consisted of 5 min of oxycodone access followed by a 25-min timeout during which both levers retracted. This 30-min trial was repeated 12 times per session for a total of 6 h per day. Under this IntA schedule, presses on the active lever resulted in a single intravenous injection of oxycodone (0.05 mg/kg, infused over 2.5 s - 4.5 s), paired with a cue light above the active lever. The dose of oxycodone was reduced (0.1 mg/kg to 0.05 mg/kg) during IntA based on our previous work indicating that reducing the dose led to robust responding during IntA sessions (Samson et al., 2022). Rats underwent one IntA self-administration session daily for ten consecutive days.

### Abstinence and cue-induced drug seeking

Following the last IntA self-administration session, rats underwent a forced abstinence period of 14 days. During this phase, rats remained in their home cage (except when performing cue-induced seeking tests). In Experiment 1, seeking tests were performed on AD1 and AD14. Due to potential extinction-like decreases in lever responding observed in a subset of rats, we conducted Experiment 2, where only one seeking test was performed on AD14. All seeking tests were 30 min in length with both active and inactivate levers available. Active lever presses resulted in a 20 s cue light presentation, but there was no oxycodone delivery. Under these conditions, the number of active lever presses served as a measure of oxycodone seeking. In both experiments, treatment with vehicle (100 nl DMSO) or suvorexant (30 mg/kg) was administered 24 h prior to the AD14 seeking test (AD13) to avoid potential sedative effects of the drug (Winrow and Renger, 2014) and in consideration of prior work from our laboratory indicating that the effects of hypocretin-based treatments are long lasting (Brodnik et al., 2020; Clark et al., 2024; Samels et al., 2024).

### Ex vivo fast-scan cyclic voltammetry

Approximately 24 h after the seeking test on AD14, rats were anesthetized with 2.5% isoflurane for 5 min and subsequently decapitated. The brain was rapidly removed and transferred to ice-cold aCSF containing NaCl (126 mM), KCl (2.5 mM), NaH2PO4 (1.2 mM), CaCl2 (2.4 mM), MgCl2 (1.2 mM), NaHCO3 (25 mM), glucose (11 mM), and L-ascorbic acid (0.4 mM). A vibratome was used to produce 400 µm-thick coronal sections containing the NAc core. Slices were transferred to room temperature oxygenated aCSF and left to equilibrate for at least 1 h before being transferred into a recording chamber with aCSF (32°C). Oxycodone-naive (naive) rats did not receive surgery and were used as a control group throughout these studies.

A bipolar stimulating electrode was placed on the surface of the tissue in the NAc core, and a carbon fiber microelectrode was implanted between the stimulating electrode leads. Dopamine release was evoked using a single electrical pulse (400 µA, 4ms, monophasic) every 3 min. Once baseline dopamine peak height was stable (3 successive stimulations within <10% variation), the slice was exposed to increasing concentrations of oxycodone (0.01, 0.1, 1, 10, and 100 µM) (Experiment 2 only). Quantification of dopamine measurements was conducted with Michaelis-Menten modelling using Demon Voltammetry and Analysis Software (Yorgason et al., 2011). Modeling of dopamine dynamics was completed using an average value for the final three collections for baseline measurements and the final three collections for each concentration of oxycodone (Experiment 2 only).

## Data Analysis

All active and inactive lever presses were recorded during behavioral sessions. Inactive lever presses served as a measure of nonspecific behavior. All rats that met acquisition criteria were used for FSCV experiments. Analyses to detect sex differences were conducted for all behavioral and voltammetry metrics. We did not observe any interactions between sex and measures of interest indicating that both females and males responded similarly across test conditions. Therefore, female and male data were combined but denoted where appropriate. For reference, Supplemental Table 1 and 2 show the results from the analyses performed to detect sex differences.

## Statistical Analysis

Statistical analyses were conducted using GraphPad Prism 9.4.0. Specific analyses are detailed in the results section. Self-administration data were analyzed using a one-way ANOVA with day as the within-subjects variable. Oxycodone seeking data were analyzed using a two-way ANOVA with seeking day as the within-subjects variable and treatment (vehicle or suvorexant) as the between-subjects variable (Experiment 1) or an unpaired t-test (Experiment 2). Baseline voltammetry measurements of DA peak height and uptake were analyzed using a one-way ANOVA with treatment (naive, vehicle or suvorexant) as the between-subjects variable. The effects of oxycodone on dopamine peak height and uptake were analyzed using a two-way ANOVA with oxycodone concentration as the within-subjects variable and treatment (naive, vehicle, and suvorexant) as the between-subjects variable.

## Results

### Suvorexant does not influence cue-induced oxycodone seeking in rats tested for incubation of oxycodone seeking

To examine the effects of suvorexant on oxycodone seeking during abstinence, rats underwent a cue-induced seeking test on AD1 and were treated with vehicle (n = 12) or 30 mg/kg suvorexant (n = 11) on AD13, 24 h prior to undergoing a second cue-induced seeking test on AD14 (**Figure 1A**). Similar to our previous report (Samson et al., 2022), rats readily acquired oxycodone self-administration (**Figure 1B**). A one-way repeated measures ANOVA revealed that IntA to oxycodone resulted in escalation of intake (*F*_(4.699, 103.4)_=3.586, *p*=0.0059) (**Figure 1C**). We found that suvorexant did not affect cue-induced seeking on AD14 (**Figure 1D**). A two-way ANOVA revealed no effect of treatment (*F*_(1,21)_=0.2675, *p*=0.6104) or seeking day (*F*_(1,21)_=0.1675, *p*=0.6865), and no treatment X seeking day interaction (*F*_(1,21)_=0.0967, *p*=0.7589) on lever presses.

**Figure 1.**
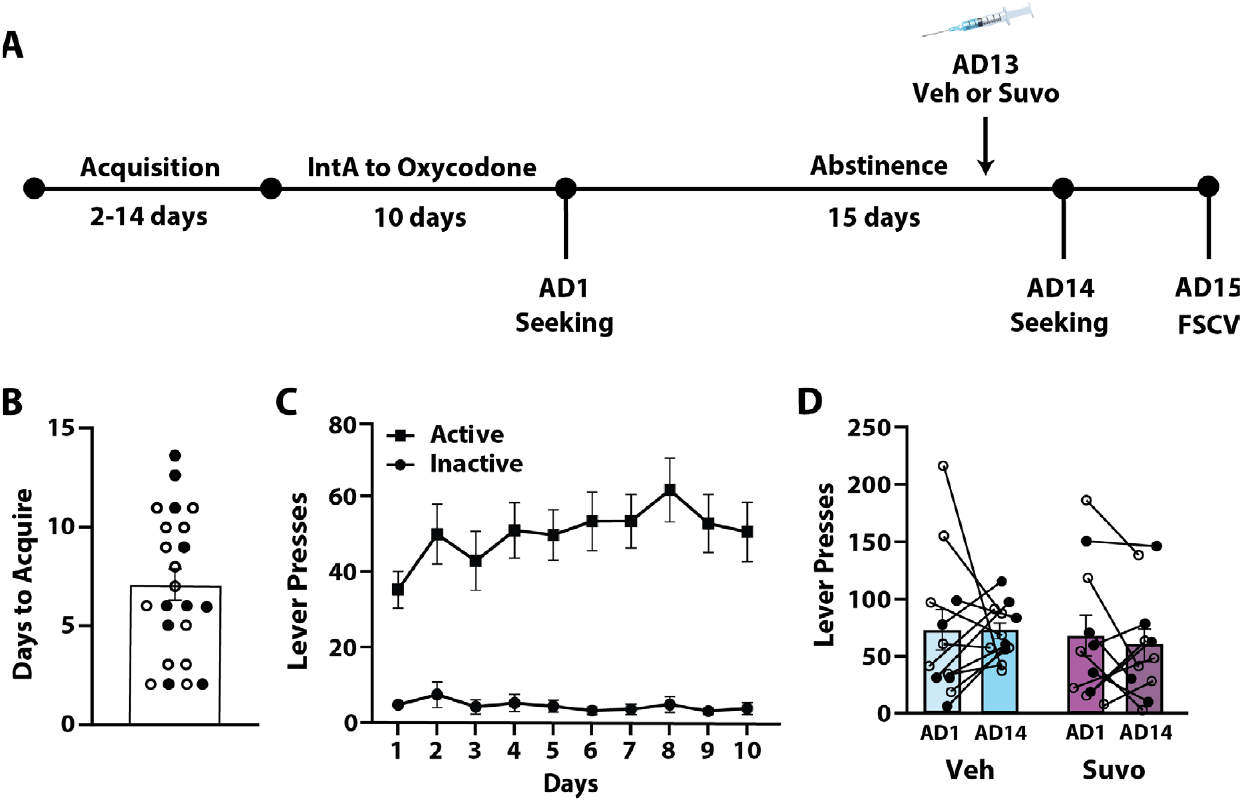
Rats exhibit escalation of oxycodone intake but not incubation of oxycodone seeking. (A) Experimental timeline. (B) Days to acquire oxycodone self-administration. (C) Active and inactive lever presses across the 10 days of IntA. (D) Lever presses during 30 min cue-induced seeking tests on abstinence day 1 (AD1) and AD14. Vehicle (Veh), n = 12 (7 females◯, 5 males•); Suvorexant (Suvo), n = 11 (6 females◯, 5 males•). Data shown are mean ± SEM.

### Dual hypocretin receptor antagonism normalized dopamine uptake in the NAc core

To examine the effects of suvorexant on dopamine transmission, a subset of rats from Experiment 1 were euthanized on AD15 for FSCV recordings in the NAc core (**Figure 1A**). A one-way ANOVA revealed no significant effect of treatment (*F*_(2,21)_=0.8070, *p*=0.6037) on dopamine peak height (**Figure 2A**). However, a one-way ANOVA revealed a significant effect of treatment (*F*_(2,21)_=4.251, *p*=0.0140) on dopamine uptake (**Figure 2B**). A Dunnett’s post-hoc tests showed that dopamine uptake was reduced in vehicle-treated rats compared to naive rats, consistent with our previous findings (Samson et al., 2022). Notably, suvorexant restored dopamine uptake, as there was no significant difference between suvorexant-treated rats and naive controls.

**Figure 2.**
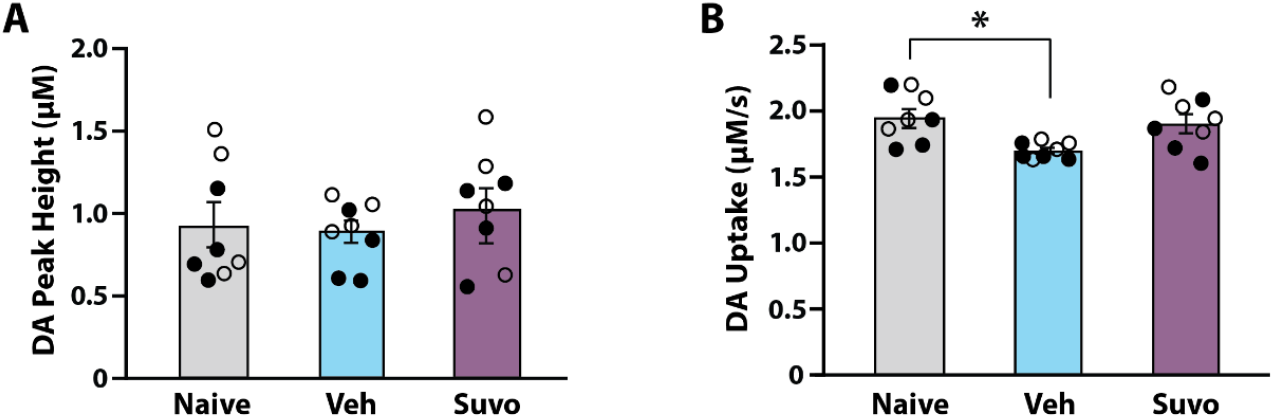
Dual hypocretin receptor antagonism normalized dopamine uptake in the NAc core. (A) Dopamine peak height and (B) dopamine uptake at baseline. Naive, n = 8 (4 females◯, 4 males•); Vehicle (Veh), n = 8 (4 females◯, 4 males•); Suvorexant (Suvo), n = 8 (4 females◯, 4 males•). Data shown are mean ± SEM. \**p*<0.05 compared to Naive.

### Suvorexant reduces oxycodone seeking in rats with sustained seeking behavior

As described above, rats subjected to seeking tests on AD1 and AD14 did not exhibit incubation of oxycodone seeking as a group, which is consistent with our previous report (Samson et al., 2022). Notably, in the current studies it appeared that a subpopulation of rats (10 out of 23) exhibited high lever pressing on AD1 (≥60 presses) and all but one of these rats reduced their lever pressing on AD14. By comparison, of the subset of rats (13 out of 23) that displayed comparatively low pressing on AD1 (<60 presses), all but two rats increased lever pressing on AD14. These observations suggest the possibility that rats with high pressing on AD1 may have experienced extinction-like behavior that could have obscured the effects of suvorexant on AD14 pressing. To circumvent the possibility of extinction, we conducted an additional experiment with a separate cohort of rats (Experiment 2) in which we omitted the AD1 seeking test, thereby removing the possibility that high levels of seeking early in abstinence could lead to extinction prior to suvorexant treatment. Rats underwent 10 days of IntA to oxycodone and then a 14-day abstinence period. On AD13, rats were treated with vehicle (n = 17) or 30 mg/kg suvorexant (n = 17) 24 h prior to the cue-induced seeking test on AD14 (**Figure 3A**). As in Experiment 1, rats readily acquired oxycodone self-administration (**Figure 3B**). Additionally, a one-way repeated measures ANOVA showed that all rats displayed escalated oxycodone intake across the 10-day IntA schedule as in Experiment 1 (*F*_(3.737, 127.1)_=6.997, *p*<0.0001) (**Figure 3C**). An unpaired Student’s t-test revealed a significant difference in lever pressing on AD14 between suvorexant and vehicle-treated rats (*t*_(33)_=0.2.660, *p*=0.0120) showing that suvorexant reduced oxycodone seeking on AD14 (**Figure 3D**).

**Figure 3.**
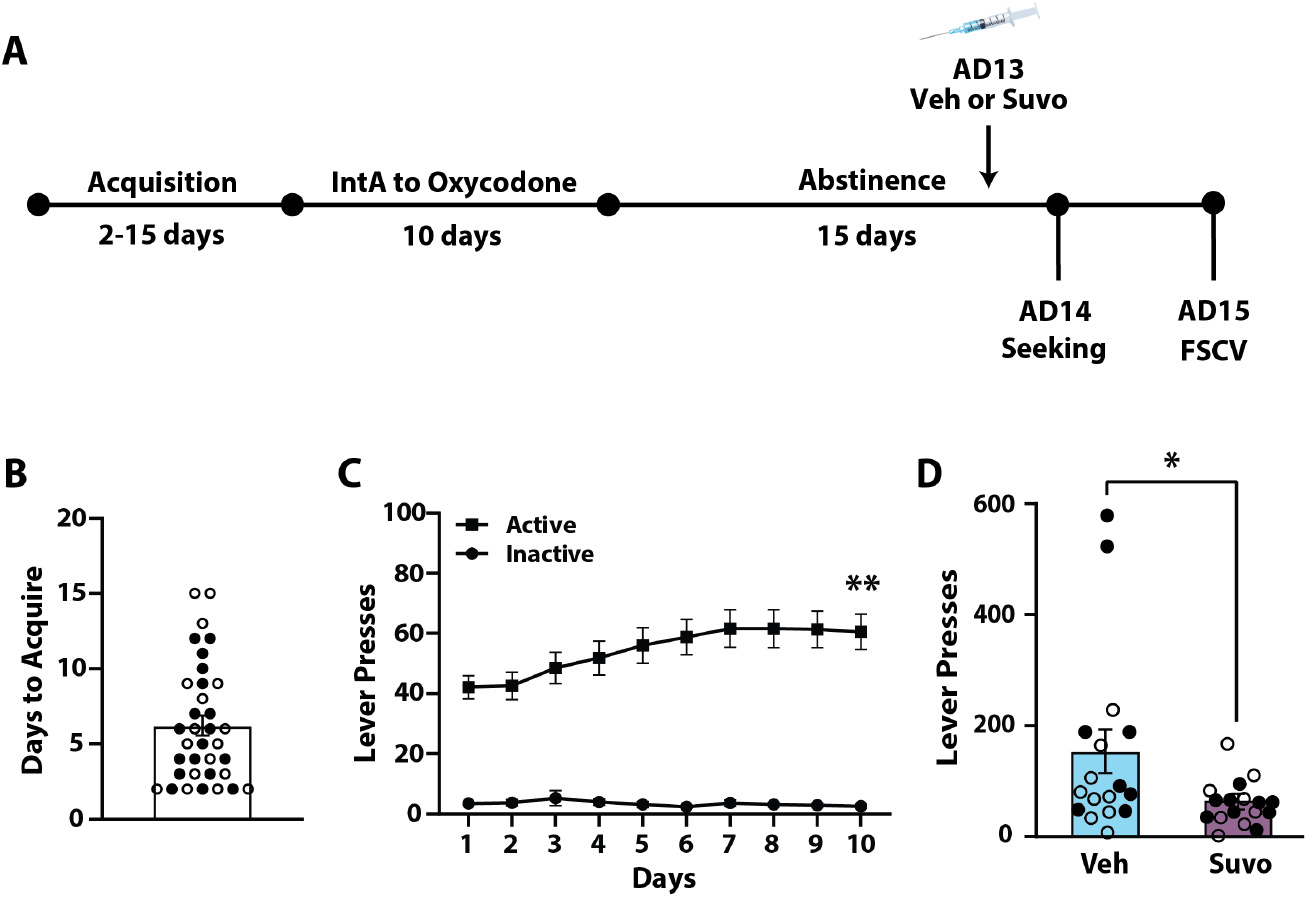
Suvorexant reduces cue-induced oxycodone seeking. (A) Experimental timeline. (B) Days to acquire self-administration for all rats. (C) Active lever presses across the 10 days of IntA for all rats and (D) Lever presses during 30 min cue-induced seeking tests on abstinence day 14 for all rats. Vehicle (Veh), n = 17 (9 females◯, 8 males•); Suvorexant (Suvo), n = 17 (8 females◯, 9 males•). Data shown are mean ± SEM. \**p*<0.05, \*\**p*<0.01 vs. Day 1 of IntA.

### Dual hypocretin receptor antagonism normalized dopamine uptake in the NAc core

To examine the effects of suvorexant on dopamine transmission, rats from Experiment 2 were used for FSCV recordings in the NAc core (**Figure 3A**). A one-way ANOVA revealed no significant effect of treatment (*F*_(2,41)_=1.542, *p*=0.3336) on dopamine peak height (**Figure 4A**). As in Experiment 1, a one-way ANOVA revealed a significant effect of treatment (*F*_(2,41)_=4.165, *p*=0.0226) on dopamine uptake (**Figure 4B**).

**Figure 4.**
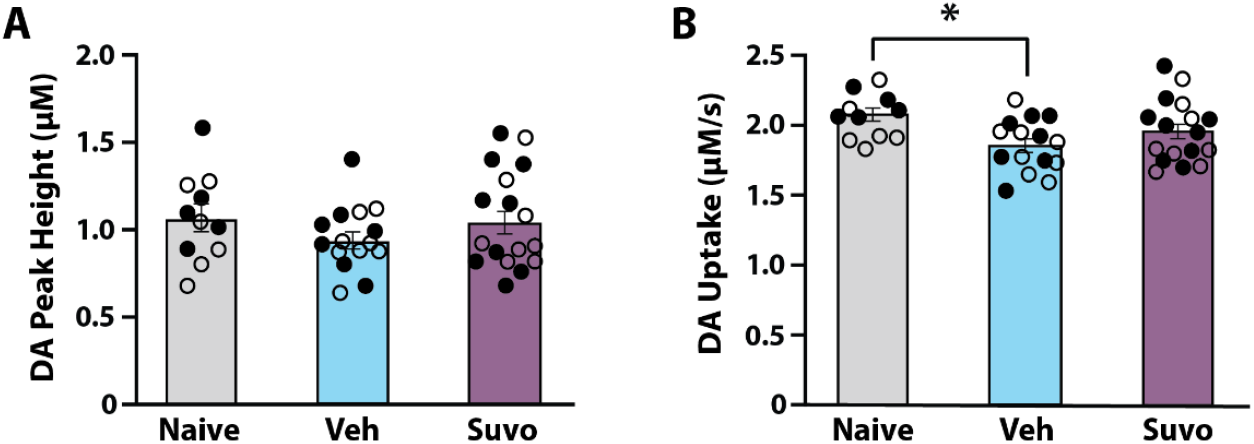
Treatment with suvorexant normalized dopamine uptake in the NAc core. (A) Dopamine peak height and (B) dopamine uptake at baseline. Naive, n = 11 (6 females◯, 5 males•); Vehicle (Veh), n = 15 (8 females◯, 7 males•); Suvorexant (Suvo), n = 17 (8 females◯, 9 males•). Data shown are mean ± SEM. \**p*<0.05 compared to Naive.

Dunnett’s post-hoc tests revealed a significant difference between vehicle-treated and naive rats, such that oxycodone exposure reduced dopamine uptake in vehicle-treated rats. This difference was not observed in suvorexant-treated rats, suggesting that suvorexant normalized dopamine uptake to some extent.

### Suvorexant did not influence the acute effects of oxycodone on dopamine peak height or uptake in the NAc core

Given our prior finding that acute oxycodone application to NAc brain slices reduced dopamine uptake (Samson et al., 2022), we examined whether suvorexant influenced the acute effects of oxycodone on dopamine transmission. We applied increasing concentrations of oxycodone to NAc slices and measured dopamine transmission in tissue from naive, vehicle-or suvorexant-treated rats from Experiment 2. A two-way ANOVA revealed a significant effect of oxycodone concentration (*F*_(1.890,66.15)_=22.96, *p*<0.0001), but no effect of treatment (*F*_(2,35)_=0.1619, *p*=0.8511) or a concentration X treatment interaction (*F*_(10,175)_=0.4974, *p*=0.8901) on dopamine peak height (**Figure 5A**). Additionally, a two-way ANOVA revealed a significant effect of oxycodone concentration (*F*_(2.193,76.77)_=179.3, *p*<0.0001), but no effect of treatment (*F*_(2,35)_=0.4311, *p*=0.6532), or a concentration X treatment interaction (*F*_(10,175)_=0.5809, *p*=0.8282) on dopamine uptake (**Figure 5B**). Consistent with our previous studies (Samson et al., 2022), oxycodone reduced dopamine uptake, but suvorexant does not differentially affect dopamine responses to acute oxycodone exposure.

**Figure 5.**
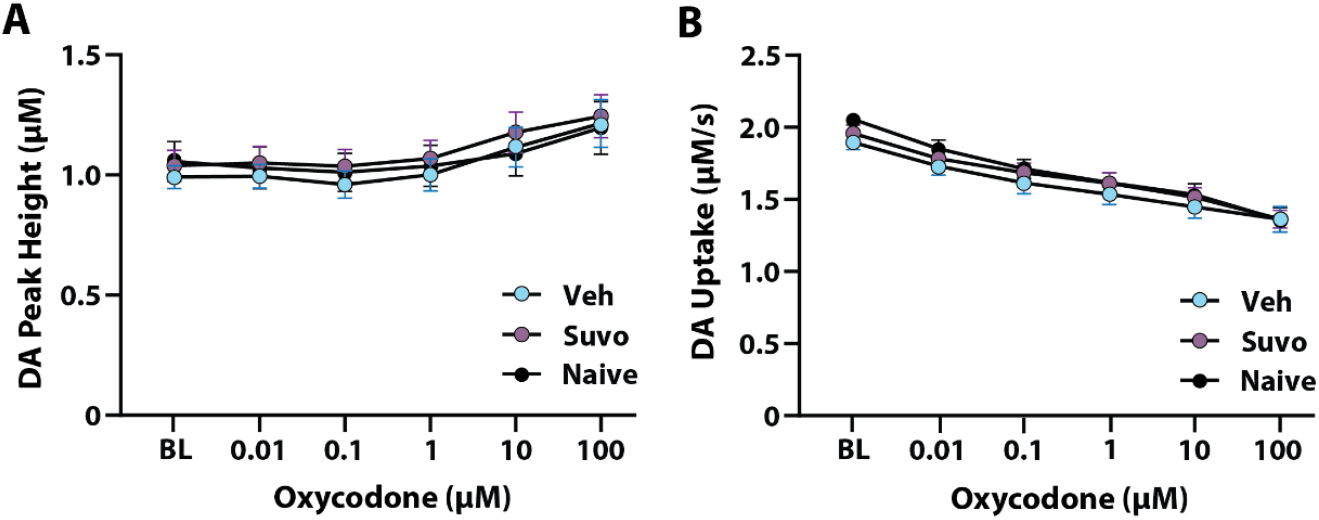
Treatment with suvorexant did not influence the acute effects of oxycodone on dopamine transmission. The effect of oxycodone on (A) dopamine peak height and (B) dopamine uptake in NAc slices. Naive, n = 10 (5 females, 5 males); Vehicle (Veh), n = 12 (7 females, 5 males); Suvorexant (Suvo), n = 16 (10 females, 6 males). Data shown are mean ± SEM.

## Discussion

In the current studies, we examined the effects of the FDA-approved dual hypocretin receptor antagonist suvorexant on oxycodone seeking and dopamine transmission during abstinence. We found that after IntA to oxycodone, suvorexant did not reduce cue-induced oxycodone seeking. However, vehicle-treated rats exhibited reduced dopamine uptake compared to naive rats, recapitulating our previous findings following IntA to oxycodone. We discovered two distinct phenotypes in these rats. Low Takers during IntA exhibited persistent oxycodone seeking both early and later in abstinence, while High Takers during IntA showed some evidence of extinction following the AD1 seeking test. Under these conditions, hypocretin receptor antagonism did not influence seeking behavior, but this was likely due to extinction observed in High Takers (Experiment 1). To avoid potential extinction effects, we omitted the AD1 seeking test and observed high lever pressing on AD14. In these studies, we found that suvorexant significantly reduced cue-induced oxycodone seeking on AD14 (Experiment 2). In line with our previous findings (Samson et al., 2022), we found that vehicle-treated rats exhibited reduced dopamine uptake and that suvorexant normalized dopamine uptake to some extent. Taken together, these observations suggest that suvorexant reduces cue-induced oxycodone seeking, likely by normalizing aberrant dopamine transmission.

### Dual hypocretin receptor antagonism reduces cue-induced oxycodone seeking

Amassing evidence indicates that the hypocretin system modulates opioid-related behaviors. For example, blockade of HcrtR1 reduces motivation for remifentanil (Porter-Stransky et al., 2017; Mohammadkhani et al., 2020), fentanyl (Fragale et al., 2019; Fragale et al., 2021), and heroin (Smith and Aston-Jones, 2012), and reduces cued reinstatement for fentanyl (Fragale et al., 2021), heroin (Smith and Aston-Jones, 2012), and oxycodone (Matzeu and Martin-Fardon, 2020). Studies assessing opioid motivation and seeking have primarily investigated the involvement of HcrtR1, with limited research exploring the contributions of HcrtR2 or the effects of dual hypocretin receptor manipulations. HcrtR2 antagonism has no effect on cued reinstatement of fentanyl seeking (Matzeu and Martin-Fardon, 2020), yet interestingly, suvorexant reduces both motivation and cued reinstatement of fentanyl seeking (O’Connor et al., 2020). Our finding that suvorexant reduces cue-induced oxycodone seeking is consistent with these observations and is the first to show that following IntA to oxycodone, hypocretin receptor antagonism reduces seeking during abstinence.

A notable aspect of our findings is that suvorexant reduced oxycodone seeking 24 h following administration. This provides evidence that hypocretin-based treatments may have behavioral effects that are long lasting. Indeed, studies have reported that HcrtR1 antagonism reduces cued reinstatement of remifentanil 24 hrs post-treatment, even though the compound is undetectable 8 hr after administration (Mohammadkhani et al., 2020). Additionally, our lab has shown that HcrtR1 antagonism reduces cocaine seeking 7 days following treatment (Clark et al., 2024; Samels et al., 2024). This evidence, which supports long-lasting behavioral effects of hypocretin manipulations, presents an intriguing prospect for future treatments for opioid use disorder. Follow up studies are needed to further clarify the duration of these effects and the mechanisms underlying the action of non-opioid, hypocretin-based treatments.

### Dual hypocretin receptor antagonism normalizes aberrant dopamine uptake following IntA to oxycodone

Extensive evidence indicates that in drug-naive rats hypocretins can bidirectionally modulate dopamine levels through actions in the VTA. For example, intra-VTA administration of hypocretin peptides increases dopamine in the NAc (Narita et al., 2006; España et al., 2011), while intra-VTA HcrtR1 antagonism (España et al., 2010) and knockdown of HcrtR1 in the VTA (Bernstein et al., 2018; Black et al., 2023) reduces dopamine in the NAc. Further, these actions of the hypocretin system on mesolimbic dopamine are thought to underlie their involvement in motivation for drugs of abuse. In support of this, knockout of the hypocretin peptide attenuates morphine-induced dopamine in the NAc (Narita et al., 2006). In the current studies, we found that IntA to oxycodone and 14 days of abstinence did not affect dopamine peak height in vehicle-or suvorexant-treated rats compared to oxycodone-naive rats (**Figure 4A**). This observation is consistent with prior reports indicating that HcrtR1, HcrtR2, or dual hypocretin receptor antagonism does not influence dopamine levels (Prince et al., 2015; Gentile et al., 2018; Brodnik et al., 2020).

Increasing evidence indicates that abstinence from opioids leads to changes in the DAT, a critical regulator of dopamine levels in the NAc. In humans, there is strong evidence of reduced DAT availability in the striatum during opioid abstinence (Shi et al., 2008; Yeh et al., 2012; Liu et al., 2013; Yuan et al., 2017; Kamp et al., 2019). Consistent with this, DAT levels are reduced in the anterior basal forebrain following chronic morphine administration in rats (Simantov, 1993). Recently, we showed that abstinence from IntA to oxycodone reduced dopamine uptake in the NAc core on both AD2 and AD15, indicating reduced efficiency of the DAT (Samson et al., 2022). This lasting reduction in dopamine uptake throughout abstinence suggests that dopamine uptake alterations may be contributing to oxycodone seeking. Therefore, we hypothesized that treatment with suvorexant, a dual hypocretin receptor antagonist, would not only reduce oxycodone seeking, but also normalize dopamine uptake. Consistent with our previous findings, we observed that vehicle-treated rats showed reduced dopamine uptake compared to naive rats (**Figure 4B**). However, suvorexant normalized dopamine uptake, suggesting that it may be reducing cue-induced oxycodone seeking through normalization of dopamine transmission.

### Sex differences in oxycodone intake under an IntA schedule

Studies exploring sex differences in opioid seeking behaviors are limited and present mixed results. For instance, several reports suggest that females acquire heroin self-administration more rapidly than males (Lynch and Carroll, 1999; Carroll et al., 2002) and that females administer more heroin than males under long access schedules (Cicero et al., 2003; Towers et al., 2019; George et al., 2021). However, other studies contradict these findings, showing no sex differences in the acquisition of heroin or fentanyl self-administration (Vazquez et al., 2020; Bakhti-Suroosh et al., 2021; George et al., 2021). Consistent with the latter studies and our previous report (Samson et al., 2022), we found no sex differences in acquisition of oxycodone self-administration although in Experiment 2 we did observe that males displayed higher intake than females. (**Supplemental Tables 1 and 2**).

Previous studies on motivation for opioids and cued reinstatement of opioid seeking have been conducted exclusively in males. Our study is the first to investigate the impact of hypocretin receptor antagonism on opioid-seeking behavior in females. Interestingly, we did not observe a significant effect of sex on suvorexant effects on oxycodone seeking (**Supplemental Table 2**). Overall, our current and prior studies contribute to the nuanced understanding of these sex-specific behaviors, underscoring the need for further research into the mechanisms underlying these differences.

## Conclusions

Substantial evidence implicates the hypocretin system in modulating dopamine transmission and motivated behaviors. We recently showed that abstinence from oxycodone reduces dopamine uptake which coincides with high levels of oxycodone seeking. In the current studies we found that treatment with suvorexant, a dual hypocretin receptor antagonist, attenuated oxycodone seeking and normalized dopamine uptake. These observations suggest that the hypocretin system may be a promising pharmaceutical target for reducing drug-associated behaviors, such as opioid craving and seeking.

## Acknowledgements

This material is based upon work supported by the National Science Foundation (NSF) Graduate Research Fellowship Program under Grant No. 2041772, the Commonwealth Universal Research Enhancement Program (CURE; RAE) and the National Institute on Drug Abuse (NIDA) grant R01DA039100 (RAE). Any opinions, findings, and conclusions or recommendations expressed in this material are those of the author(s) and do not necessarily reflect the views of the NSF, CURE, or NIDA.

## Author Contributions

KRS, conceptualized and designed the studies, conducted the self-administration and FSCV data collection, analyzed and interpreted the data, wrote the original draft, and edited the content. ARB, conducted the self-administration and FSCV data collection, wrote and edited the content. RAE, conceptualized and designed the studies, analyzed and interpreted the data, edited the original draft, and edited the content.

## References

Bakhti-Suroosh A, Towers EB, Lynch WJ (2021) A buprenorphine-validated rat model of opioid use disorder optimized to study sex differences in vulnerability to relapse. Psychopharmacology 238:1029–1046.

Bernosky-Smith KA, Stanger DB, Trujillo AJ, Mitchell LR, España RA, Bass CE (2016) The GLP-1 agonist exendin-4 attenuates self-administration of sweetened fat on fixed and progressive ratio schedules of reinforcement in rats. Pharmacology Biochemistry and Behavior 142:48–55.

Bernstein DL, Badve PS, Barson JR, Bass CE, España RA (2018) Hypocretin receptor 1 knockdown in the ventral tegmental area attenuates mesolimbic dopamine signaling and reduces motivation for cocaine. Addiction biology 23:1032–1045.

Black EM, Samels SB, Xu W, Barson JR, Bass CE, Kortagere S, España RA (2023) Hypocretin / Orexin Receptor 1 Knockdown in GABA or Dopamine Neurons in the Ventral Tegmental Area Differentially Impact Mesolimbic Dopamine and Motivation for Cocaine. Addict Neurosci 7.

Blackwood CA, Leary M, Salisbury A, McCoy MT, Cadet JL (2019a) Escalated oxycodone self-administration causes differential striatal mRNA expression of FGFs and IEGs following abstinence-associated incubation of oxycodone craving. Neuroscience 415:173–183.

Blackwood CA, Hoerle R, Leary M, Schroeder J, Job MO, McCoy MT, Ladenheim B, Jayanthi S, Cadet JL (2019b) Molecular adaptations in the rat dorsal striatum and hippocampus following abstinence-induced incubation of drug seeking after escalated oxycodone self-administration. Molecular neurobiology 56:3603–3615.

Brodnik ZD, Alonso IP, Xu W, Zhang Y, Kortagere S, España RA (2020) Hypocretin receptor 1 involvement in cocaine-associated behavior: Therapeutic potential and novel mechanistic insights. Brain Res 1731:145894.

Carroll ME, Morgan AD, Lynch WJ, Campbell UC, Dess NK (2002) Intravenous cocaine and heroin self-administration in rats selectively bred for differential saccharin intake: phenotype and sex differences. Psychopharmacology (Berl) 161:304–313.

Cicero TJ, Aylward SC, Meyer ER (2003) Gender differences in the intravenous self-administration of mu opiate agonists. Pharmacology Biochemistry and Behavior 74:541–549.

Clark PJ, Migovich VM, Das S, Xi W, Kortagere S, España RA (2024) Hypocretin Receptor 1 Blockade Early in Abstinence Prevents Incubation of Cocaine Seeking and Normalizes Dopamine Transmission. bioRxiv:2024.2011.2030.625912.

España RA, Melchior JR, Roberts D, Jones SR (2011) Hypocretin 1/orexin A in the ventral tegmental area enhances dopamine responses to cocaine and promotes cocaine self-administration. Psychopharmacology 214:415–426.

España RA, Oleson EB, Locke JL, Brookshire BR, Roberts DC, Jones SR (2010) The hypocretin–orexin system regulates cocaine self-administration via actions on the mesolimbic dopamine system. European Journal of Neuroscience 31:336–348.

Fragale JE, James MH, Aston-Jones G (2021) Intermittent self-administration of fentanyl induces a multifaceted addiction state associated with persistent changes in the orexin system. Addiction biology 26:e12946.

Fragale JE, Pantazis CB, James MH, Aston-Jones G (2019) The role of orexin-1 receptor signaling in demand for the opioid fentanyl. Neuropsychopharmacology 44:1690–1697.

Gentile TA, Simmons SJ, Barker DJ, Shaw JK, España RA, Muschamp JW (2018) Suvorexant, an orexin/hypocretin receptor antagonist, attenuates motivational and hedonic properties of cocaine. Addiction biology 23:247–255.

George BE, Barth SH, Kuiper LB, Holleran KM, Lacy RT, Raab-Graham KF, Jones SR (2021) Enhanced heroin self-administration and distinct dopamine adaptations in female rats. Neuropsychopharmacology 46:1724–1733.

Illenberger JM, Flores-Ramirez FJ, Matzeu A, Mason BJ, Martin-Fardon R (2023) Suvorexant, an FDA-approved dual orexin receptor antagonist, reduces oxycodone self-administration and conditioned reinstatement in male and female rats. Frontiers in Pharmacology 14.

Kamp F, Proebstl L, Penzel N, Adorjan K, Ilankovic A, Pogarell O, Koller G, Soyka M, Falkai P, Koutsouleris N (2019) Effects of sedative drug use on the dopamine system: a systematic review and meta-analysis of in vivo neuroimaging studies. Neuropsychopharmacology 44:660–667.

Liu Y, Han M, Liu X, Deng Y, Li Y, Yuan J, Lv R, Wang Y, Zhang G, Gao J (2013) Dopamine transporter availability in heroin-dependent subjects and controls: longitudinal changes during abstinence and the effects of Jitai tablets treatment. Psychopharmacology 230:235–244.

Lynch WJ, Carroll ME (1999) Sex differences in the acquisition of intravenously self-administered cocaine and heroin in rats. Psychopharmacology (Berl) 144:77–82.

Matzeu A, Martin-Fardon R (2020) Targeting the orexin system for prescription opioid use disorder: Orexin-1 receptor blockade prevents oxycodone taking and seeking in rats. Neuropharmacology 164:107906.

Mohammadkhani A, James MH, Pantazis CB, Aston-Jones G (2020) Persistent effects of the orexin-1 receptor antagonist SB-334867 on motivation for the fast acting opioid remifentanil. Brain research 1731:146461.

Narita M, Nagumo Y, Hashimoto S, Narita M, Khotib J, Miyatake M, Sakurai T, Yanagisawa M, Nakamachi T, Shioda S (2006) Direct involvement of orexinergic systems in the activation of the mesolimbic dopamine pathway and related behaviors induced by morphine. Journal of Neuroscience 26:398–405.

O’Connor SL, Fragale JE, James MH, Aston-Jones G (2020) The dual orexin/hypocretin receptor antagonist suvorexant reduces addiction-like behaviors for the opioid fentanyl. bioRxiv:2020.2004.2025.061887.

Porter-Stransky KA, Bentzley BS, Aston-Jones G (2017) Individual differences in orexin-I receptor modulation of motivation for the opioid remifentanil. Addiction biology 22:303–317.

Prince CD, Rau AR, Yorgason JT, España RA (2015) Hypocretin/Orexin regulation of dopamine signaling and cocaine self-administration is mediated predominantly by hypocretin receptor 1. ACS chemical neuroscience 6:138–146.

Rasmussen K, White DA, Acri JB (2019) NIDA’s medication development priorities in response to the Opioid Crisis: ten most wanted. Neuropsychopharmacology 44:657–659.

Samels SB, Shaw JK, Alonso P, Black EM, España RA (2024) Hypocretin receptor 1 blockade early in abstinence reduces future demand for cocaine. bioRxiv:2024.2012.2006.627226.

Samson KR, Xu W, Kortagere S, España RA (2022) Intermittent access to oxycodone decreases dopamine uptake in the nucleus accumbens core during abstinence. Addict Biol 27:e13241.

Schmeichel BE, Barbier E, Misra KK, Contet C, Schlosburg JE, Grigoriadis D, Williams JP, Karlsson C, Pitcairn C, Heilig M (2015) Hypocretin receptor 2 antagonism dose-dependently reduces escalated heroin self-administration in rats. Neuropsychopharmacology 40:1123–1129.

Shi J, Zhao L-Y, Copersino ML, Fang Y-X, Chen Y, Tian J, Deng Y, Shuai Y, Jin J, Lu L (2008) PET imaging of dopamine transporter and drug craving during methadone maintenance treatment and after prolonged abstinence in heroin users. European journal of pharmacology 579:160–166.

Simantov R (1993) Chronic morphine alters dopamine transporter density in the rat brain: possible role in the mechanism of drug addiction. Neuroscience letters 163:121–124.

Simmons SJ, Martorana R, Philogene-Khalid H, Tran FH, Gentile TA, Xu X, Su S, Rawls SM, Muschamp JW (2017) Role of hypocretin/orexin receptor blockade on drug-taking and ultrasonic vocalizations (USVs) associated with low-effort self-administration of cathinone-derived 3,4-methylenedioxypyrovalerone (MDPV) in rats. Psychopharmacology (Berl) 234:3207–3215.

Smith RJ, Aston-Jones G (2012) Orexin/hypocretin 1 receptor antagonist reduces heroin self-administration and cue-induced heroin seeking. European Journal of Neuroscience 35:798–804.

Thompson AC, Zapata A, Justice JB, Jr., Vaughan RA, Sharpe LG, Shippenberg TS (2000) Kappa-opioid receptor activation modifies dopamine uptake in the nucleus accumbens and opposes the effects of cocaine. J Neurosci 20:9333–9340.

Towers EB, Tunstall BJ, McCracken ML, Vendruscolo LF, Koob GF (2019) Male and female mice develop escalation of heroin intake and dependence following extended access. Neuropharmacology 151:189–194.

Vafaie N, Kober H (2022) Association of Drug Cues and Craving With Drug Use and Relapse: A Systematic Review and Meta-analysis. JAMA psychiatry.

Vazquez M, Frazier JH, Reichel CM, Peters J (2020) Acute ovarian hormone treatment in freely cycling female rats regulates distinct aspects of heroin seeking. Learn Mem 27:6–11.

Winrow C, Renger J (2014) Discovery and development of orexin receptor antagonists as therapeutics for insomnia. British journal of pharmacology 171:283–293.

Xiao Z-w, Cao C-y, Wang Z-x, Li J-x, Liao H-y, Zhang X-x (2006) Changes of dopamine transporter function in striatum during acute morphine addiction and its abstinence in rhesus monkey. Chinese medical journal 119:1802–1807.

Yeh TL, Chen KC, Lin S-H, Lee IH, Chen PS, Yao WJ, Lee S-Y, Yang YK, Lu R-B, Liao M-H (2012) Availability of dopamine and serotonin transporters in opioid-dependent users—a two-isotope SPECT study. Psychopharmacology 220:55–64.

Yorgason JT, España RA, Jones SR (2011) Demon voltammetry and analysis software: analysis of cocaine-induced alterations in dopamine signaling using multiple kinetic measures. J Neurosci Methods 202:158–164.

Yuan J, Liu XD, Han M, Lv RB, Wang YK, Zhang GM, Li Y (2017) Comparison of striatal dopamine transporter levels in chronic heroin-dependent and methamphetamine-dependent subjects. Addiction biology 22:229–234.

Zimmer BA, Oleson EB, Roberts D (2012) The motivation to self-administer is increased after a history of spiking brain levels of cocaine. Neuropsychopharmacology 37:1901–1910.

