## Supplemental Tables for "Dual Hypocretin Receptor Antagonism Reduces Oxycodone Seeking During Abstinence"

**Supplemental Table 1:**

| **Experiment 1** | | | |
|---|---|---|---|
| **Measure** | **Dependent Variable** | **Result** | **Test** |
| **Days to Acquire** | Sex | $t_{(21)}$=0.4443, $p$=0.6614 | Student's t-test |
| **Oxycodone Intake during IntA** | Day of IntA (1-10)<br>Sex<br>Day of IntA X Sex | $F_{(1.429, 30.01)}$=3.013, $p$=0.0794<br>$F_{(1,21)}$=4.700, $p$=0.0418*<br>$F_{(9,189)}$=2.474, $p$=0.1763 | Two-Way ANOVA |
| **Effect of Suvorexant on Oxycodone Seeking** | Treatment<br>Seeking Day (AD1, AD14)<br>Sex<br>Treatment X Seeking Day<br>Treatment X Seeking Day X Sex | $F_{(1,19)}$=0.2991, $p$=0.5908<br>$F_{(1,19)}$=0.0377, $p$=0.8481<br>$F_{(1,19)}$=0.0134, $p$=0.9092<br>$F_{(1,19)}$=0.1756, $p$=0.6799<br>$F_{(1,19)}$=1.202, $p$=0.2867 | Three-Way ANOVA |
| **Effect of Suvorexant on Dopamine Peak Height** | Treatment<br>Sex<br>Treatment X Sex | $F_{(1,12)}$=1.267, $p$=0.2824<br>$F_{(1,12)}$=2.354, $p$=0.1509<br>$F_{(1,12)}$=0.0188, $p$=0.8931 | Two-Way ANOVA |
| **Effect of Suvorexant on Dopamine Uptake** | Treatment<br>Sex<br>Treatment X Sex | $F_{(2,18)}$=5.902, $p$=0.0107*<br>$F_{(1,18)}$=2.522, $p$=0.1297<br>$F_{(2,18)}$=0.4340, $p$=0.6545 | Two-Way ANOVA |

**Supplemental Table 2:**

| **Experiment 2** | | | |
|---|---|---|---|
| **Measure** | **Dependent Variable** | **Result** | **Test** |
| **Days to Acquire** | Sex | $t_{(33)}$=0.2651, $p$=0.7926 | Student's t-test |
| **Oxycodone Intake during IntA** | Day of IntA (1-10)<br>Sex<br>Day of IntA X Sex | $F_{(3.635, 105.4)}$=7.104, $p$<0.0001*<br>$F_{(1,33)}$=7.419, $p$=0.0102*<br>$F_{(9,296)}$=1.041, $p$=0.4074 | Two-Way ANOVA |
| **Effect of Suvorexant on Oxycodone Seeking** | Treatment<br>Sex<br>Treatment X Sex | $F_{(1,30)}$=6.616, $p$=0.0153*<br>$F_{(1,30)}$=2.387, $p$=0.1329<br>$F_{(1,30)}$=3.395), $p$=0.0753 | Two-Way ANOVA |
| **Effect of Suvorexant on Dopamine Peak Height** | Treatment<br>Sex<br>Treatment X Sex | $F_{(2,38)}$=1.094, $p$=0.3453<br>$F_{(1,38)}$=1.662, $p$=0.2051<br>$F_{(2,38)}$=0.2189, $p$=0.8044 | Two-Way ANOVA |
| **Effect of Suvorexant on Dopamine Uptake** | Treatment<br>Sex<br>Treatment X Sex | $F_{(2,38)}$=4.228, $p$=0.0220*<br>$F_{(1,38)}$=0.1898, $p$=0.1898<br>$F_{(2,38)}$=0.2443), $p$=0.7845 | Two-Way ANOVA |
| **Effect of Suvorexant on Acute Effects of Oxycodone on Dopamine Peak Height** | Oxycodone Concentration<br>Treatment<br>Sex | $F_{(1.861,59.54)}$=21.33, $p$<0.001*<br>$F_{(2,32)}$=0.113, $p$=0.894<br>$F_{(1,32)}$=0.534, $p$=0.47 | Three-Way ANOVA |
| **Effect of Suvorexant on Acute Effects of Oxycodone on Dopamine Uptake** | Oxycodone Concentration<br>Treatment<br>Sex | $F_{(2.226,71.23)}$=166.240, p<0.001*<br>$F_{(2,32)}$=0.381, $p$=0.686<br>$F_{(1,32)}$=0.548, $p$=0.465 | Three-Way ANOVA |
